# Bacterial Community Structure and Diversity of Common Mosquito Species in Chengdu: Insights from PacBio Third-Generation

**DOI:** 10.1101/2025.05.27.656271

**Authors:** Xing Yifan, Lu Rong, Tian Wenjia, Li Zelin, Zhang Wei, Xu Kai, Deng Liangli, Fan Shuangfeng

## Abstract

Mosquitoes, as critical vectors of diseases such as Japanese encephalitis, dengue fever, and yellow fever, pose significant public health risks in Chengdu, a subtropical city in southwestern China. This study integrated ecological surveillance and PacBio third-generation sequencing to characterize the symbiotic microbiota of four dominant mosquito species (Aedes albopictus, Culex pipiens, Culex tritaeniorhynchus, and Armigeres subalbatus) across urban and rural habitats. From 2020 to 2024, mosquito density monitoring revealed spatial heterogeneity(Aedes albopictus, Culex pipiens, Culex tritaeniorhynchus, and Anopheles sinensis), with outer ring areas exhibiting the highest density (34.69 mosquitoes per trap), while central urban zones had the lowest (3.60). Sequencing identified 717 high-quality Amplicon Sequence Variants (ASVs), with Aedes albopictus harboring the most unique bacterial species (191). Beta diversity analysis demonstrated distinct microbial clustering among species, driven by Pseudomonadota dominance (54.27–93.89%) and variations in secondary phyla (Bacteroidota, Campylobacterota). Functional prediction via KEGG highlighted elevated human disease-associated pathways in Ae. albopictus, contrasting with reduced environmental adaptation activity. Notably, Wolbachia (clade B) and Klebsiella variicola exhibited species-specific abundance patterns, underscoring their roles in pathogen suppression and public health risks. Unclassified taxa (norank_d Bacteria, Candidatus_Hydrogenedentes) clustered near novel mosquito-associated spirochetes, suggesting underexplored functional microbiota. This study provides foundational data for understanding mosquito-microbe interactions and informs strategies for mitigating vector-borne disease.

**Author Summary:** Mosquito-borne diseases such as dengue fever and Japanese encephalitis pose severe public health risks in subtropical regions, yet critical gaps remain in understanding how mosquito-associated microbes influence disease transmission in under-resourced areas. Chengdu, a megacity in southwestern China, faces unique challenges due to rapid urbanization, climatic suitability for mosquito proliferation, and disparities in public health infrastructure compared to eastern coastal regions. This study integrates five years of ecological surveillance (2020–2024) and PacBio third-generation sequencing to map mosquito density patterns and characterize symbiotic bacterial communities in four dominant mosquito species (Aedes albopictus, Culex pipiens, Culex tritaeniorhynchus, and Armigeres subalbatus).

Key findings reveal stark spatial heterogeneity in mosquito density, with rural outer-ring areas harboring 10-fold higher densities (34.69 mosquitoes/trap) than urban centers. Notably, Aedes albopictus exhibited the highest diversity of unique symbiotic bacteria (191 species), including Wolbachia (clade B), known to suppress arboviruses like dengue. Conversely, Armigeres subalbatus carried high abundances of Klebsiella variicola, an emerging human pathogen linked to severe infections. Functional analyses further highlighted elevated human disease-associated pathways in Ae. albopictus, underscoring its dual role as a disease vector and microbial reservoir.

This work provides the first comprehensive baseline data on mosquito-microbe interactions in Chengdu, identifying actionable targets for biocontrol (e.g., leveraging Wolbachia) and early-warning systems for pathogen surveillance. By bridging ecological, molecular, and public health perspectives, our findings offer critical insights for mitigating neglected tropical diseases in subtropical regions where socioeconomic disparities intersect with high disease burdens.

## Introduction

As a critical medical vector, mosquitoes transmit a spectrum of diseases, including Japanese encephalitis, dengue fever, and yellow fever, resulting in approximately 1 million deaths and over 700 million infections annually[1]. Chengdu, situated in the western Sichuan Basin, is characterized by a subtropical humid climate with synchronized rainy and hot seasons, short frost periods, and abundant green vegetation, creating an ideal habitat for mosquito proliferation. In recent years, ecological shifts, climate variability, and the hosting of large-scale international events have exacerbated the risks of emerging, re-emerging, and imported mosquito-borne diseases, imposing substantial pressure on local public health systems. Given the strong correlation between mosquito population distribution and disease transmission dynamics, there is an urgent need to investigate local mosquito density patterns to refine current prevention and control strategies.

Concurrently, advancements in research on insect-associated symbiotic microbiota have deepened our understanding of host-microbe interactions. Numerous studies demonstrate that symbiotic bacteria regulate critical physiological processes in insect hosts, including immune defense, reproductive modulation, and developmental pathways [2–5]. Specifically, certain mosquito symbionts influence vector competence for pathogen transmission [6, 7], while gut microbiota enhance insecticide resistance by upregulating detoxification enzymes and stimulating metabolic pathways[8].

Elucidating and leveraging these symbiotic microorganisms for mosquito biocontrol represents an innovative frontier in combating vector-borne diseases.

With advancements in sequencing technologies, Academician Xu Jianguo proposed the “reverse pathogenomics” framework in 2019, a paradigm distinct from Koch’s postulates. This approach emphasizes proactive discovery of novel microorganisms in vectors via sequencing, followed by rapid assessment of their pathogenic potential and public health implications, enabling early deployment of targeted strategies to preempt emerging epidemics [9]. This methodology circumvents limitations inherent to traditional culture-based workflows (isolation, cultivation, and biochemical identification), which are often impractical for low-abundance or fastidious bacteria in small vectors like mosquitoes—particularly those requiring specialized conditions (e.g., anaerobic, low-temperature) and involving labor-intensive, time-consuming, and subjective biochemical analyses.

To address these challenges, this study integrated mosquito light trap surveillance with third-generation high-throughput sequencing (PacBio) of the 16S rRNA gene. We conducted baseline monitoring of dominant wild mosquito species in Chengdu,to map their spatiotemporal density variations. Concurrently, we characterized the structure and diversity of their symbiotic microbiota. By identifying functional microbial resources, this work provides critical insights into the coevolutionary dynamics and interaction mechanisms between mosquitoes and their symbionts. Furthermore, it facilitates early detection of uncharacterized or exotic microorganisms, enabling the establishment of a pathogen surveillance framework. These efforts aim to inform proactive strategies for controlling endemic mosquito populations and mitigating risks posed by emerging, re-emerging, or unknown vector-borne diseases in Chengdu.

## 1. Materials and methods

### 1.1 Ecological Monitoring of Adult Mosquitoes

From 2020 to 2024, adult mosquito surveillance was conducted in 22 districts of Chengdu (excluding Dongbu New Area) using the mosquito light trap method. In urban areas, two sites each were selected in residential zones, parks, and hospitals. In rural areas, two sites each were chosen in residential houses and livestock sheds. All monitoring sites, except livestock sheds, were located in outdoor environments. Surveillance was performed twice monthly from April to November each year, with a minimum interval of 10 days between consecutive sessions. Two mosquito traps were deployed per habitat. Traps were activated one hour before sunset and deactivated one hour after sunrise the following day. Captured mosquitoes were morphologically identified to species level based on characteristics such as the mesonotal scutum, tarsal segments, and color patterns, using standardized taxonomic keys.

### 1.2 Sampling of Mosquito Specimens for Symbiotic Bacterial Sequencing

Based on preliminary surveillance data, four national mosquito surveillance sites were selected in Dujiangyan City, Pengzhou City, Chongzhou City, and Qingyang District of Chengdu. Live mosquito specimens were collected using human-baited traps from July to September 2024..Collected specimens were transported to the laboratory on the same day. Morphological identification of mosquito species was performed under stereomicroscopy according to taxonomic keys, focusing on diagnostic characteristics such as mesoscutum patterns, tarsal segment morphology, and pigmentation bands. Post-identification, blood-fed females were excluded from the sample pool. All confirmed specimens were cryopreserved at −80°C for subsequent molecular analyses.

### 1.3 DNA extraction and PCR amplification

Total microbial genomic DNA was extracted from mosquito samples using the FastPure Stool DNA Isolation Kit (MJYH, shanghai, China). The DNA extract was checked on 1% agarose gel, and DNA concentration and purity were determined with NanoDrop2000 spectrophotometer (Thermo Scientific, United States).the bacterial 16S rRNA genes were amplified using the universal bacterial primers 27F (5’-AGRGTTYGATYMTGGCTCAG-3’) and 1492R (5’-RGYTACCTTGTTACGACTT-3’). Primers were tailed with PacBio barcode sequences to distinguish each sample. Amplification reactions (20-μL volume) consisted of 2×Pro Taq 10 μL, forward primer (5 μM) 0.8 μL, reverse primer (5 μM) 0.8 μL, template DNA 10 ng and DNase-free water. The PCR amplification was performed as follows: initial denaturation at 95 °C for 3 min, followed by 30 cycles of denaturing at 95 °C for 30 s, annealing at 60 °C for 30 s and extension at 72 °Cfor 45 s, and single extension at 72 °C for 10 min, and end at 4 °C (T100 Thermal Cycler PCR thermocycler, BIO-RAD, USA). After electrophoresis, The PCR products were purified using the AMPure® PB beads (Pacifc Biosciences, CA, USA) and quantified with Qubit 4.0 (Thermo Fisher Scientific, USA).

### 1.4 DNA library construction and sequencing

Purified products were pooled in equimolar and DNA library was constructed using the SMRTbell prep kit 3.0 (Pacifc Biosciences, CA, USA) according to PacBio’s instructions. Purified SMRTbell libraries were sequenced on the Pacbio Sequel IIe System (Pacifc Biosciences, CA, USA) by Majorbio Bio-Pharm Technology Co. Ltd. (Shanghai, China). High-fidelity (HiFi) reads were obtained from the subreads, generated using circular consensus sequencing via SMRT Link v11.0.

### 1.5 Data processing and statistical analysis

HiFi reads were barcode-identified and length-filtered. For bacterial 16S rRNA gene, sequences with a length <1,000 or >1,800 bp were removed. The optimized-HiFi reads were de-noised using DADA2 plugin in the Qiime2[10] (version 2020.2) pipeline with recommended parameters, which obtains single nucleotide resolution based on error profiles within samples. DADA2 denoised sequences are usually called amplicon sequence variants (ASVs). To minimize the effects of sequencing depth on alpha and beta diversity measure, the number of sequence from each sample was rarefied to 6,000, which still yielded an average Good’s coverage of 97.90%. Taxonomic assignment of ASVs was performed using the Blast consensus taxonomy classifier implemented in Qiime2 and the SILVA 16S rRNA database (v138). The metagenomic function was predicted by PICRUSt2 (Phylogenetic Investigation of Communities by Reconstruction of Unobserved States) based on ASV representative sequences. PICRUSt2[11] is a software containing a series of tools as follows: HMMER was used to aligns ASV representative sequences with reference sequences. EPA-NG and Gappa were used to put ASV representative sequences into a reference tree. The castor was used to normalize the 16S gene copies. MinPath was used to predict gene family profiles, and locate into the gene pathways. Entire analysis process was accord to protocols of PICRUSt2.

### 1.6 Statistical Analysis

Bioinformatic analysis of the mosquito symbiotic microbiota was carried out using the Majorbio Cloud platform (https://cloud.majorbio.com). Based on the ASVs information, rarefaction curves and alpha diversity indices including observed ASVs, Chao1 richness, Shannon index and Good’s coverage were calculated with Mothur v1.30.1[12]. The similarity among the microbial communities in different samples was determined by principal coordinate analysis (PCoA) based on Bray-curtis dissimilarity using Vegan v2.5-3 package. The PERMANOVA test was used to assess the percentage of variation explained by the treatment along with its statistical significance using Vegan v2.5-3 package. The linear discriminant analysis (LDA) effect size (LEfSe) (http://huttenhower.sph.harvard.edu/LEfSe) was performed to identify the significantly abundant taxa (phylum to genera) of bacteria among the different groups (LDA score > 2, P < 0.05).

### 1.7 Analysis of Ecological Surveillance Data

Raw mosquito surveillance data were organized, analyzed, and visualized using WPS software. For spatial stratification, six central districts (Gaoxin, Wuhou, Chenghua, Jinjiang, Qingyang, Jinniu) were classified as the central urban area, characterized by high population density and urbanization levels. Eleven peripheral districts/counties (Dujiangyan, Pengzhou, Qingbaijiang, Jintang, Jianyang, Xinjin, Chongzhou, Dayi, Qionglai, Pujiang) were designated as the outer ring areas, dominated by mountainous and rural habitats. Regions between the central and outer rings (Wenjiang, Pidu, Xindu, Longquanyi, Shuangliu, Tianfu New Area) were categorized as the secondary ring areas, representing transitional ecological zones.

Statistical analyses were performed using SPSS 18.0. One-way ANOVA under a completely randomized design was applied to assess differences in mosquito density across years, months, and spatial strata. Pairwise comparisons of multiple sample means were conducted using the Bonferroni correction, with a significance threshold of P < 0.05.

## 2. Results

### 2.1 Mosquito Density Surveillance Results

From 2020 to 2024, a total of 9,581 mosquito traps were deployed in Chengdu, capturing 195,967 mosquitoes. The density indices (mosquitoes per trap per night) for key species were as follows: Culex pipiens (7.41),Culex tritaeniorhynchus (8.83), Anopheles sinensis (3.80), Aedes albopictus (0.41), and Aedes aegypti (0). Annual average mosquito densities from 2020 to 2024 were 13.56, 16.25, 9.32, 34.86, and 27.87, respectively, with no statistically significant interannual differences (F= 1.366,P= 0.266). Seasonal trends exhibited a unimodal distribution, peaking in June (99.67) and July (87.10). Monthly variations in density were statistically significant (F = 4.153,P= 0.002).

Spatially, mosquito density varied significantly across regions (F= 8.354,P< 0.001): the outer ring areas showed the highest density (34.69), followed by secondary ring areas (12.29), and the central urban areas had the lowest density (3.60),Fig1A. Species-specific spatial patterns were observed:Culex pipiens predominated in urbanized regions (primary and secondary rings), while Culex tritaeniorhynchus and Anopheles sinensis dominated in rural outer ring areas. For detailed spatial and temporal distributions, Fig1B.

**Fig 1.**
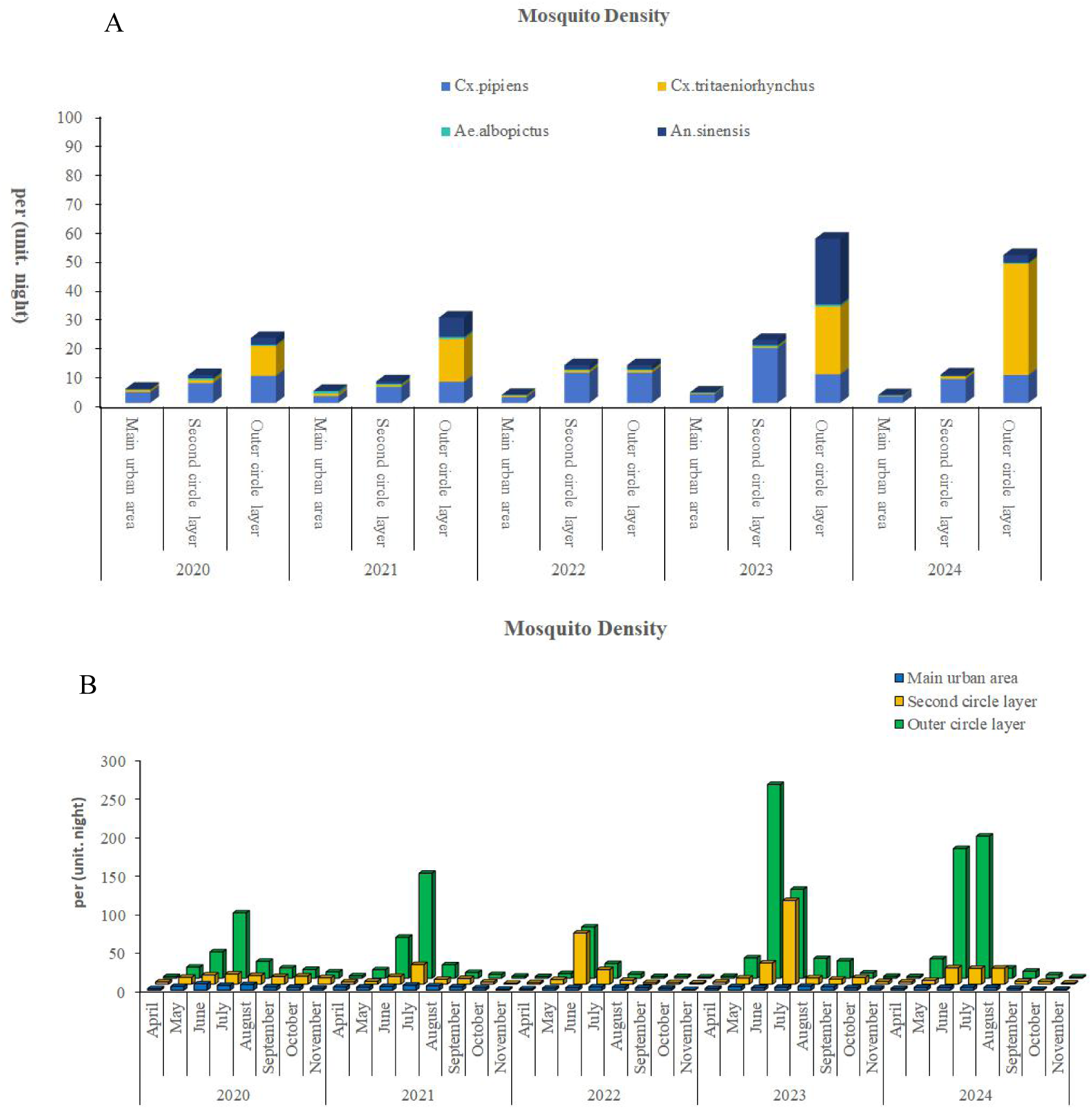
Mosquito Density Surveillance Results. Note:(A) Temporal variation in mosquito density in Chengdu, 2020–2024; (B) Changes in species-specific distribution of mosquitoes in Chengdu, 2020–2024.

### 2.2 Sequencing Results and ASV Classification

A total of 297,649 high-quality sequences were obtained after quality control, with an average sequence length of 1,490 bp (range: 1,111–1,782 bp). Sequences were clustered at 100% similarity and rarefied to the minimum sequence length (4,090 bp) to standardize sequencing depth. Taxonomic annotation of Amplicon Sequence Variants (ASVs) was performed using QIIME2, yielding 717 ASVs corresponding to 436 bacterial species. Venn diagram analysis of ASV abundance revealed distinct microbial community structures among mosquito species (Fig 2). Aedes albopictus exhibited the highest number of unique symbiotic bacterial species (191), while Culex tritaeniorhynchus harbored the fewest unique species (19). Only four bacterial species were shared across all four mosquito species.

**Fig 2.**
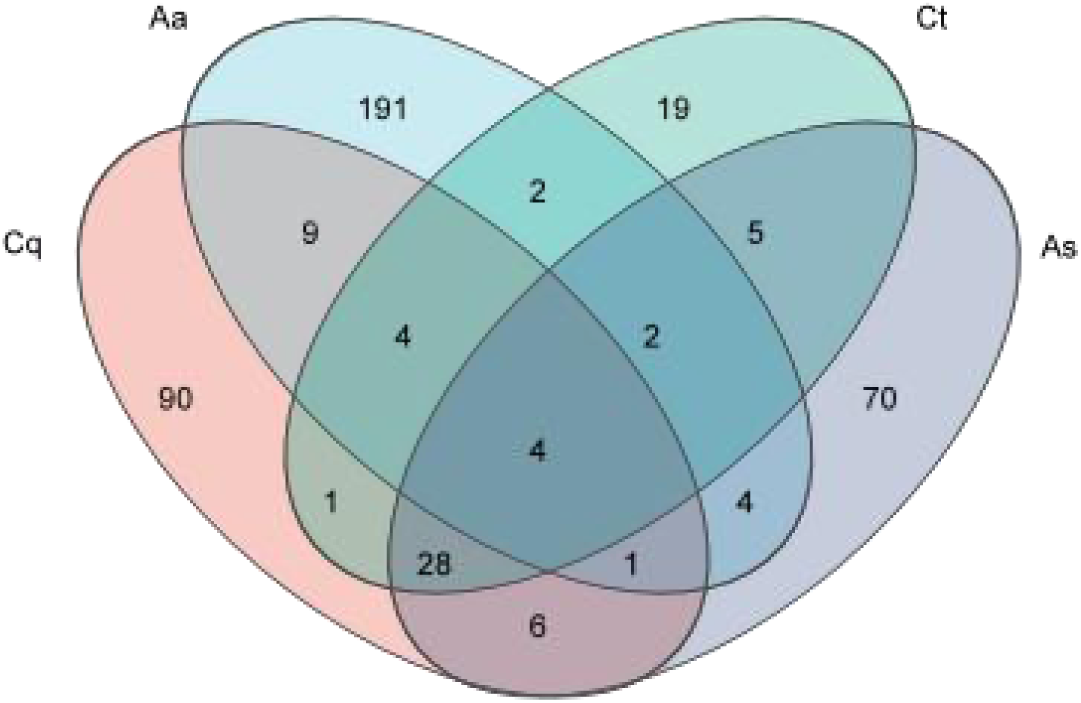
Venn diagram of symbiotic bacteria.

### 2.3 Alpha Diversity Analysis

Rarefaction curves (Fig 3) were generated by randomly subsampling sequencing data to assess sequencing depth saturation. When subsampling exceeded 500 sequences, Shannon index curves for all samples plateaued, indicating sufficient sequencing depth to capture the majority of microbial species. Alpha diversity indices (Simpson, Shannon, Chao1, and coverage) were statistically analyzed to evaluate microbial diversity and species richness(Fig4). Chao and ACE indices exhibited positive correlations with total community richness, while the Shannon index positively correlated with diversity and the Simpson index inversely correlated with diversity. The Aedes albopictus group (Aa) displayed the highest mean values for ACE (59.32 ± 23.76), Chao1 (58.75 ± 23.87), and Sobs (58.5 ± 23.90), whereas the

**Fig 3.**
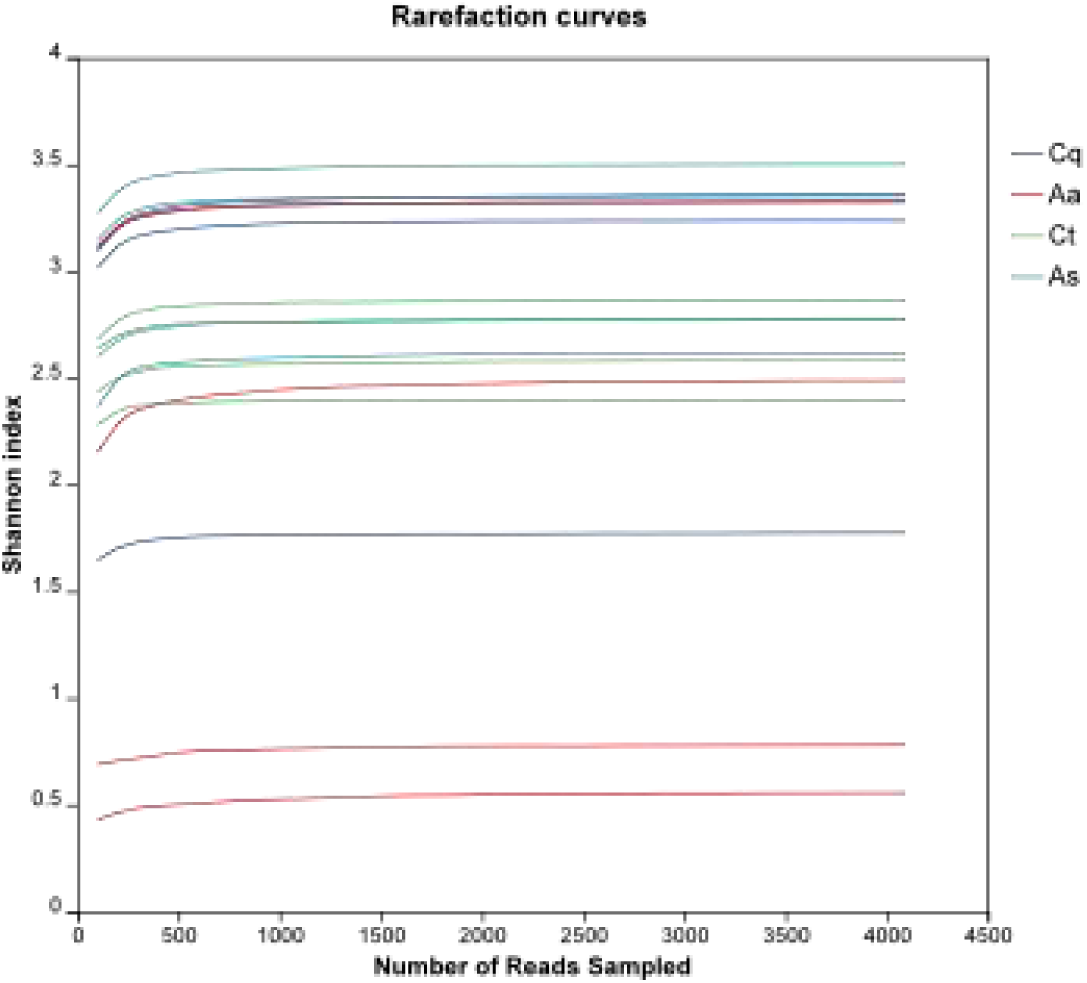
Rarefaction curves in different mosquito species. Note:**Cq**: *Culex pipiens*; **Aa**: *Aedes albopictus*;**Ct**: *Culex tritaeniorhynchus*;**As**: *Aedes pullatus* :

**Fig 4.**
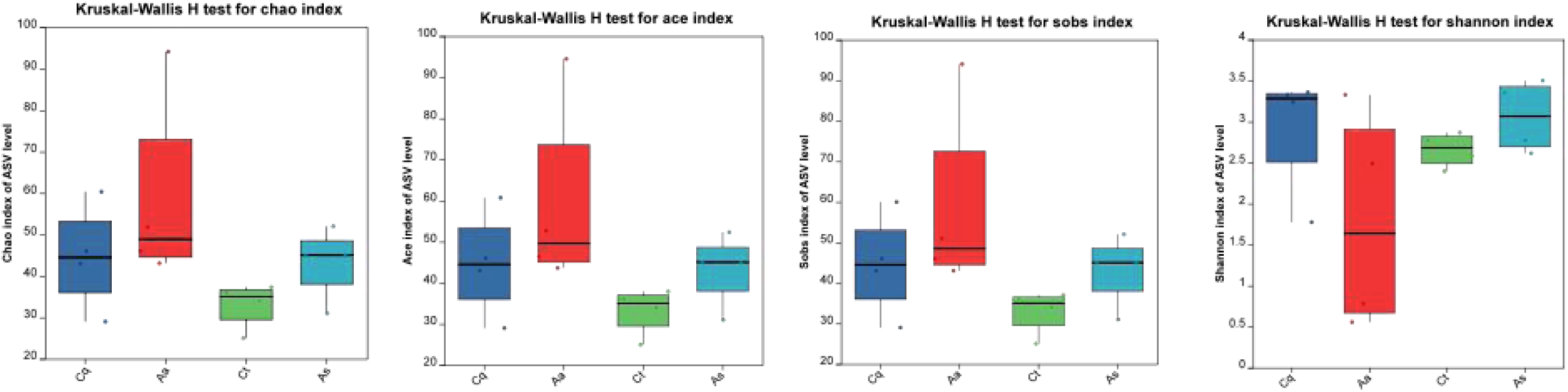
Distribution of Alpha diversity index boxplot. Note:**Cq**: *Culex pipiens*; **Aa**: *Aedes albopictus*;**Ct**: *Culex tritaeniorhynchus*;**As**: *Aedes pullatu*

Culex tritaeniorhynchus group (Ct) exhibited the lowest values (ACE: 33.22 ± 5.71; Chao: 33.08 ± 5.56; Sobs: 33 ± 5.48). The Armigeres subalbatus group (As) demonstrated the highest Shannon index (3.06 ± 0.43) and the lowest Simpson index (0.098 ± 0.082). Wilcoxon rank-sum tests revealed no statistically significant differences in richness, diversity, or evenness among mosquito species after multiple-test correction (Padjust > 0.05).

### 2.4 Beta Diversity Analysis

Beta diversity was assessed using Weighted UniFrac distances and visualized via Principal Coordinate Analysis (PCoA) with ANOSIM (999 permutations). The PCoA plot (Figure 5) illustrated distinct clustering patterns among mosquito species, where proximity along the X- and Y-axes (relative distance units) indicated higher microbial community similarity. Ellipses represent 95% confidence intervals.Culex tritaeniorhynchus(Ct) and Armigeres subalbatus (As) exhibited partial overlap, while Aedes albopictus (Aa) remained distinctly separated from other groups. ANOSIM yielded an R-value of 0.43 (P= 0.001), indicating moderate differentiation (0.25 ≤ R < 0.5) in symbiotic microbiota composition among species.

**Fig 5:**
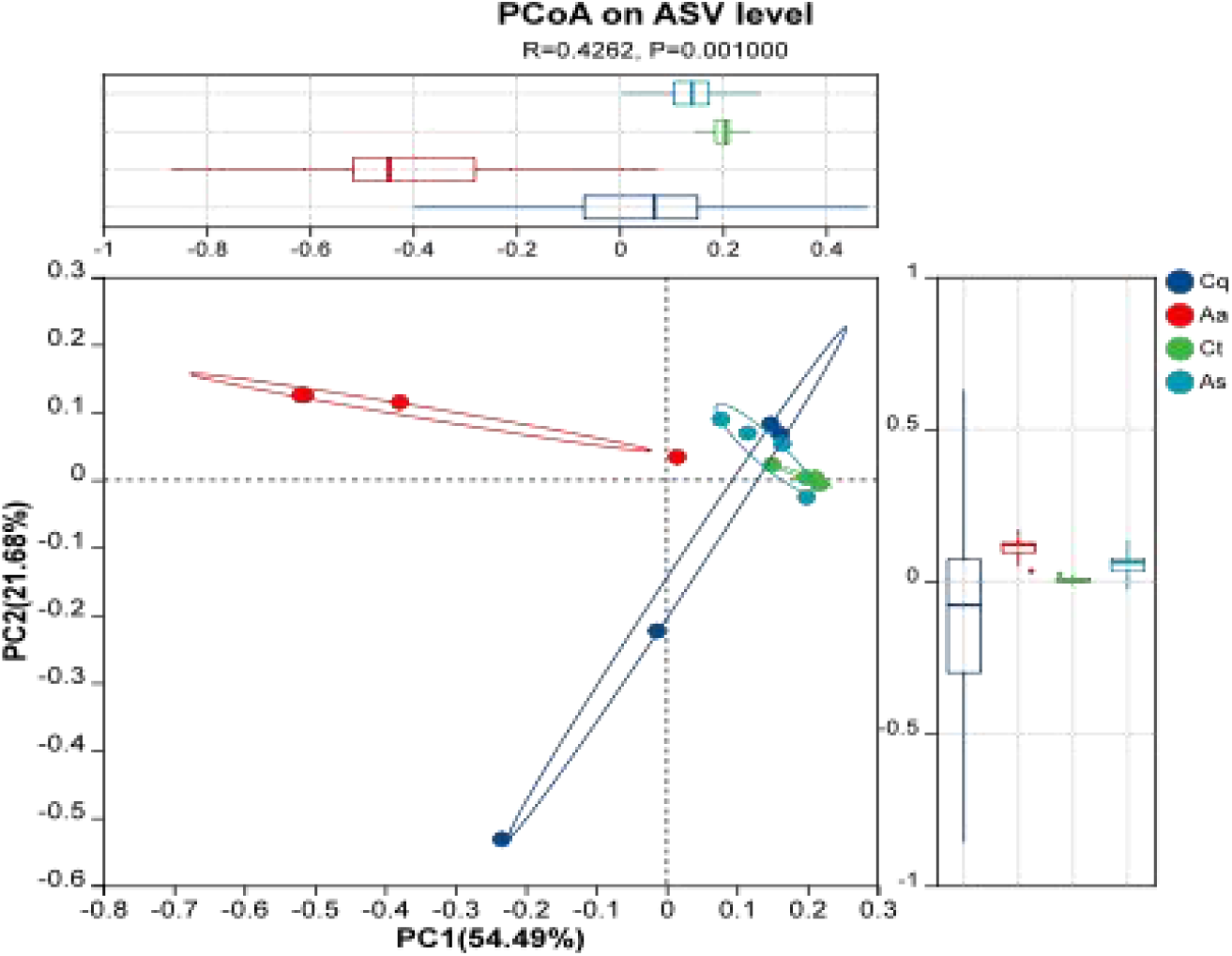
Principal Coordinates Analysis (PCoA) based on Weighted UniFrac distance.

### 2.5 UPGMA Clustering Analysis

Hierarchical clustering (UPGMA algorithm) and Bray-Curtis distance-based heatmap analysis (Figure 6) quantified interspecific microbial abundance differences. Branch lengths in the dendrogram reflected phylogenetic distances, while heatmap color gradients (red = greater divergence) highlighted sample dissimilarity. Aedes albopictus (A-a) formed a distinct clade, whereas Culex pipiens quinquefasciatus(Cp) and Culex tritaeniorhynchus(Ct) clustered closely, suggesting genus-level compositional similarity. Notably, one Armigeres subalbatus subgroup(As4) exhibited proximity to Culex tritaeniorhynchus (Ct3), diverging from conspecific subgroups, aligning with PCoA results.

**Fig 6:**
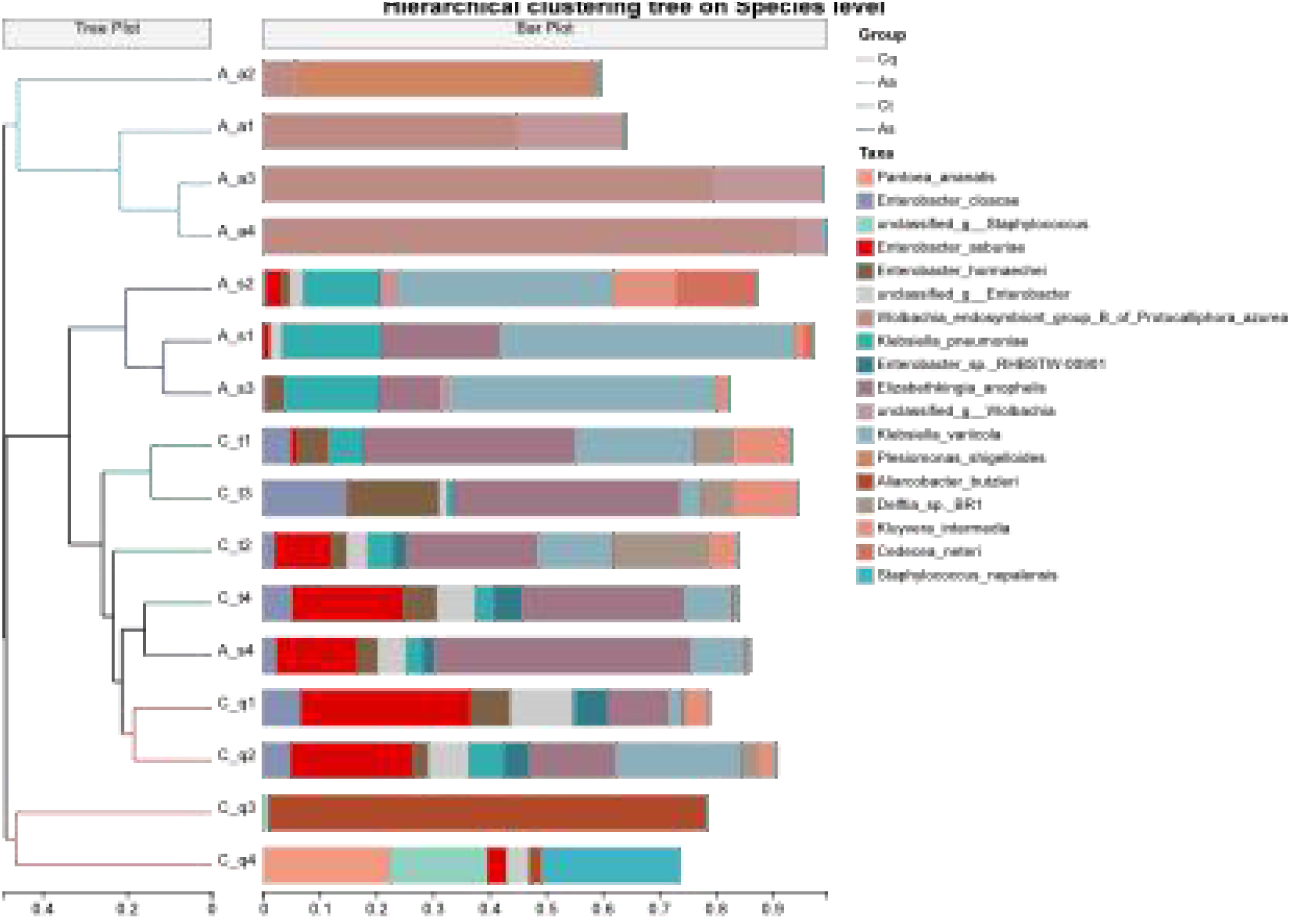
Hierarchical clustering tree on Species level.

### 2.6. Microbial Community Structure and Differential Analysis

Taxonomic annotation identified 23 phyla, 38 classes, 60 orders, 84 families, 116 genera, and 179 species (Figure 7). Dominant phyla included Pseudomonadota (relative abundance: 54.27% in Cp, 93.89% in Aa, 63.15% in Ct, 79.81% in As), Bacteroidota (36.57% in Ct, 19.59% in As), Campylobacterota (19.96% in Cp), and Fusobacteriota (3.37% in Aa). At the genus level, Enterobacter (28.03%), Aliarcobacter (19.96%), and Staphylococcus (16.66%) dominated Cp; Wolbachia (66.90%), Plesiomonas (13.03%), and Acinetobacter (4.36%) prevailed in Aa;

**Figure 7.**
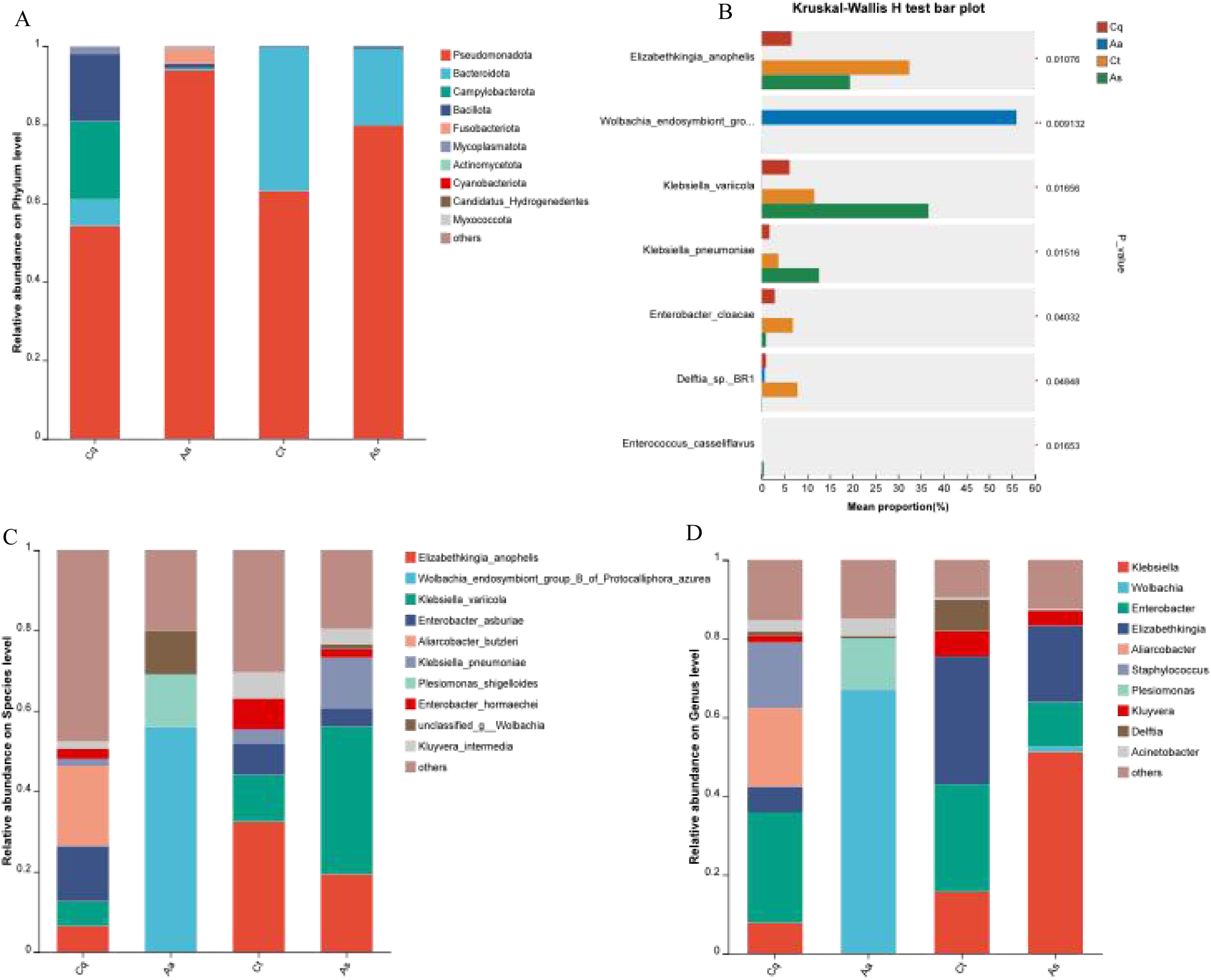
Differential analysis of species relative abundance at different taxonomic levels. **Note:A.** Relative abundance of bacterial communities in different mosquito species at the phylum level.**B.** Differential analysis of species abundance (bar chart).**C.** Relative abundance of bacterial communities in different mosquito species at the species level.**D.** Relative abundance of bacterial communities in different mosquito species at the genus level.

Elizabethkingia (32.46%), Enterobacter (27.24%), and Klebsiella (15.70%) were prominent in Ct; and Klebsiella (51.27%),Elizabethkingia(19.43%), and Enterobacter (19.43%) dominated As (Figure 6). Species-level analysis revealed *Aliarcobacter butzleri* (19.96%, Cp), *Wolbachia endosymbiont group B* of *Protocalliphora azurea* (55.98%, Aa), *Elizabethkingia anophelis* (32.46%, Ct), and *Klebsiella variicola*(36.67%, As) as dominant species.

At the species level, interspecies differences in bacterial abundance among mosquito groups were analyzed using the Kruskal-Wallis rank-sum test, with results adjusted for multiple comparisons via the false discovery rate (FDR) method. Significant differences (P < 0.05) were observed in the following species:Elizabethkingia anophelis (P= 0.011),Wolbachia endosymbiont group B of Protocalliphora azurea(P= 0.009), Klebsiella variicola(P= 0.016), Klebsiella pneumoniae(P = 0.015) Enterobacter cloacae (P= 0.04),Enterococcus casseliflavus (P = 0.017),Delftia sp. BR1 (P= 0.048),Significance levels:**0.001 < P ≤ 0.01; *0.01 < P ≤ 0.05.

### 2.7 Microbial species abundance clustering analysis

At the genus level, Taxonomic annotation and abundance profiling were conducted across two dimensions: mosquito species groups and individual samples. A heatmap was generated with the horizontal axis representing sample or group labels, the vertical axis listing microbial species names, and a phylogenetic clustering tree at the family level displayed on the left. Color gradients in the heatmap visually represented species distribution across samples, with warmer hues indicating higher abundance values.

As illustrated in Fig 8A, distinct clustering patterns of symbiotic bacteria were observed among the four mosquito species. Aedes albopictus and Armigeres subalbatus each contained three taxa with relatively high abundance (values > 1,000), whereas Culex pipiens and Culex tritaeniorhynchus exhibited four and seven high-abundance taxa, respectively. Average linkage clustering of individual samples (Fig 8B) further demonstrated moderate variations in microbial abundance among subgroups within the same mosquito species.

**Fig 8.**
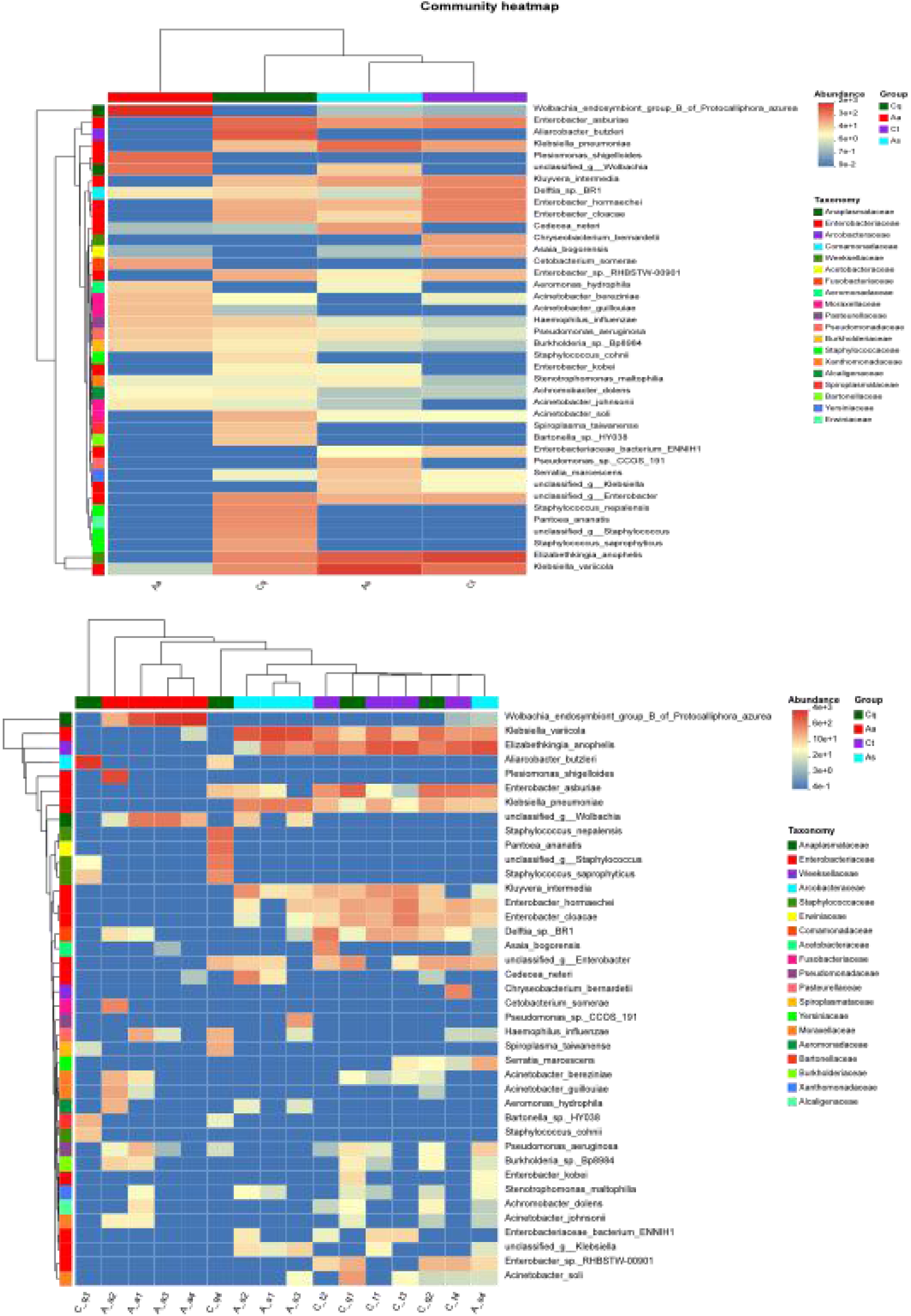
Species Abundance Clustering Heatmap. **Note: (A)** Intra-sample species abundance clustering heatmap; **(B)** Inter-species (mosquito) abundance clustering heatmap.

### 2.8 Phylogenetic Analysis of Bacterial Species

A maximum likelihood phylogenetic tree was constructed based on all Amplicon Sequence Variants (ASVs) using representative sequences annotated against the SILVA database (Fig 9). At the phylum level,Pseudomonadota was ubiquitously distributed across all four mosquito species, dominating in Aedes albopictus and Culex tritaeniorhynchus. In contrast, Bacteroidota exhibited preferential enrichment in Armigeres subalbatus and Culex pipiens, likely associated with species-specific ecological niches or physiological requirements.Unclassified taxa, such as *norank_f Enterobacteriaceae* and *norank_p Bacteroidota*, clustered within core evolutionary clades of Pseudomonadota and Bacteroidota, respectively, despite lacking genus-level classification. These taxa showed high phylogenetic affinity to known genera (Kluyvera and Sphingobacterium), suggesting their potential membership within these taxonomic groups. Notably, *norank_p Candidatus_Hydrogenedentes*, the sole representative of the candidate phylum Hydrogenedentes, formed a basal branch in the tree and clustered with *norank_d Bacteria*. Both taxa displayed close phylogenetic proximity to Entomospira nematocera, a recently identified mosquito-associated spirochete, indicating the presence of undercharacterized functional microbiota within this lineage.

**Fig 9.**
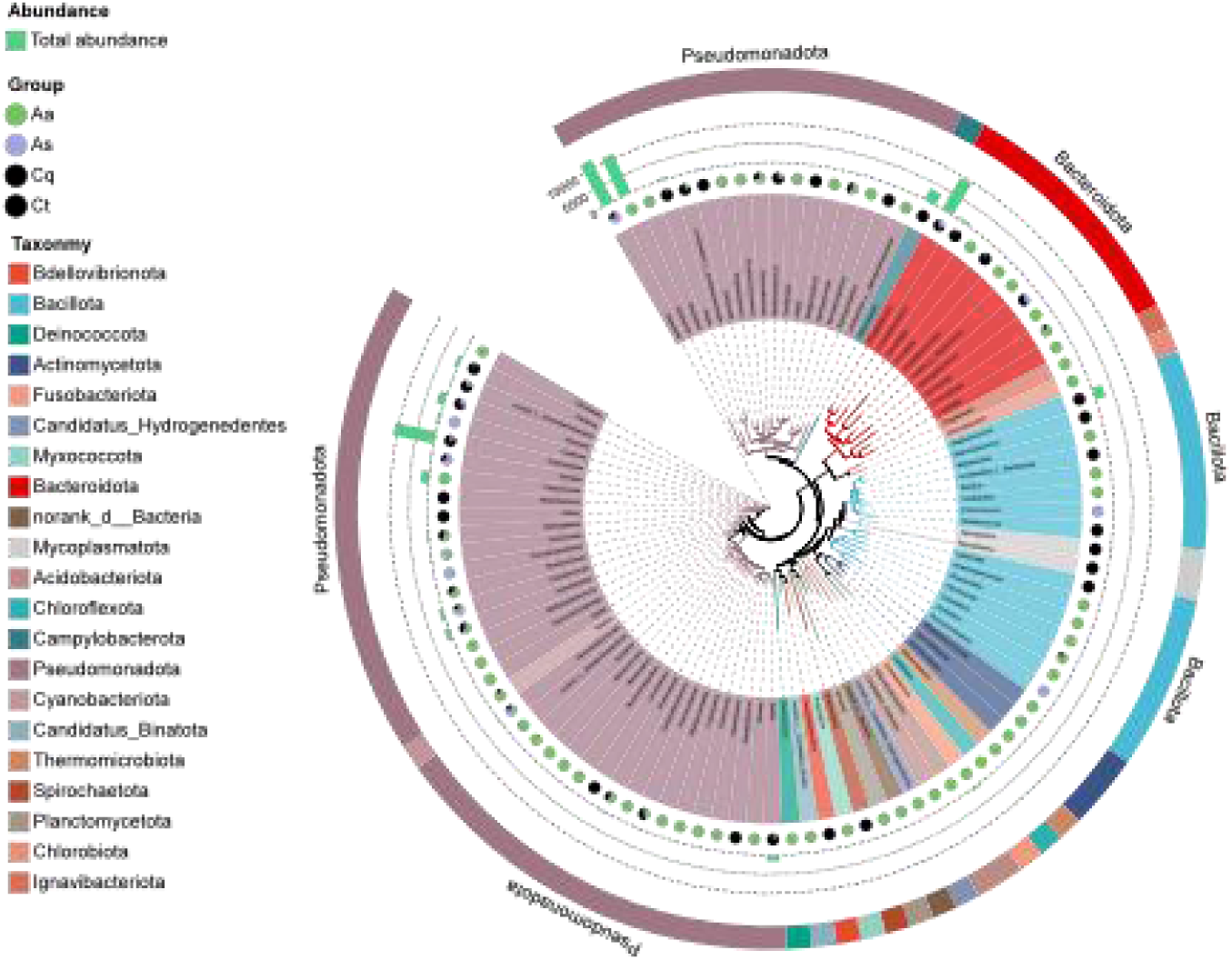
Species-level phylogenetic tree. **Note:** Pie segment colors indicate corresponding phyla. Branches represent distinct bacterial species. Stacked bar charts and proportional pie charts illustrate the abundance distribution of these bacteria across different mosquito species.

### 2.9 Functional Prediction of Microbial Communities

Functional annotation of metabolic pathways in mosquito-associated symbiotic bacteria was performed using PICRUSt2 (v2.2.0-b) based on the Kyoto Encyclopedia of Genes and Genomes (KEGG). A total of 334 functional pathways were identified, and a heatmap was generated to visualize the top 30 most abundant pathways (Fig 10). The heatmap displays functional abundance distributions across mosquito species using color gradients, where the horizontal axis represents group labels, the vertical axis lists pathway names, and color intensity is proportional to abundance.

**Fig 10.**
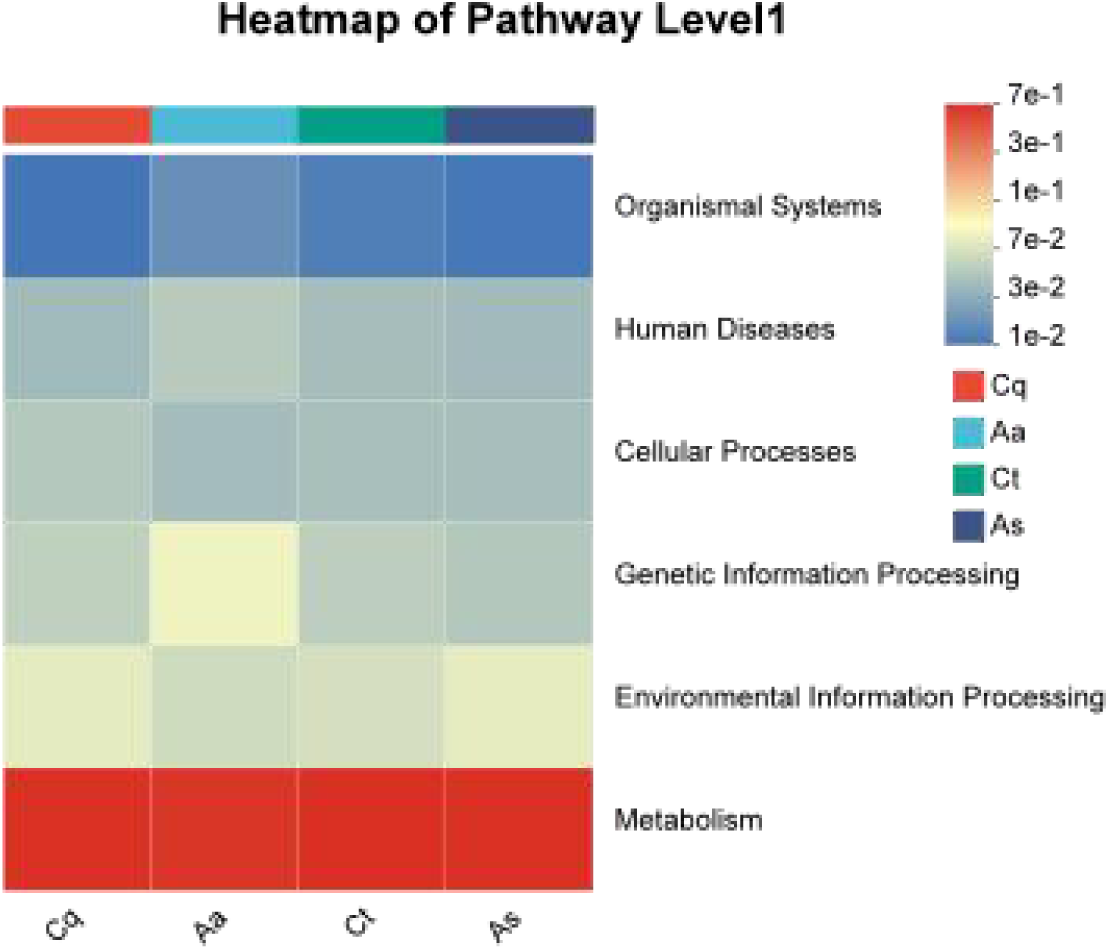
KEGG functional abundance prediction analysis heatmap Discussion.

As shown in Fig 9, significant differences in pathway abundances were observed among groups, particularly in categories such as human diseases, genetic information processing, and environmental adaptation. Compared to the other three groups, Aedes albopictus exhibited higher relative abundances in human disease-associated and genetic information processing pathways. In contrast, pathways related to cellular processes and environmental information processing were markedly less enriched in Aedes albopictus.

Mosquitoes pose a significant global health burden by transmitting pathogens responsible for diseases such as Japanese encephalitis, malaria, and dengue fever, threatening approximately 3.9 billion people across 120 countries [13]. Chengdu, a subtropical city in southwestern China, provides an ideal habitat for mosquito proliferation due to its climatic conditions and urbanization. As a hub for tourism, trade, and international events, Chengdu faces heightened risks of emerging or re-emerging mosquito-borne diseases. Historical surveillance data (2013–2017) identified Culex pipiens (50.43%), Culex tritaeniorhynchus (35.85%), and Armigeres subalbatus (7.39%) as the dominant mosquito species in Chengdu. Our analysis of 2020–2024 surveillance data revealed no significant interannual differences in overall mosquito density. However, spatial stratification showed the highest density in outer ring areas (34.69), followed by secondary ring areas (12.29), and the lowest in central urban zones (3.60). This spatial pattern likely reflects variations in vegetation cover, ecological diversity, human mobility, and public health intervention efficacy. Additionally, the rural predominance of Cx. Tritaeniorhynchus and Anopheles sinensis aligns with previous studies.

Our sequencing results revealed striking interspecies differences in symbiotic microbiota composition. Aedes albopictus harbored the highest number of unique bacterial species (191), while only four species were shared across all four mosquito groups. Although alpha diversity metrics (richness, Shannon index) showed numerical variations among species, no statistically significant differences were observed. In contrast, beta diversity analyses (UPGMA, PCoA) demonstrated distinct clustering patterns: Culex tritaeniorhynchus and Culex pipiens exhibited closer phylogenetic proximity, with partial overlap to Armigeres subalbatus, whereas Aedes albopictus formed a separate cluster. These findings align with prior studies highlighting niche-driven microbial divergence in mosquitoes [14]. At the phylum level, Pseudomonadota dominated all species (54.27–93.89%), while secondary dominant phyla varied (Bacteroidota in Cx. tritaeniorhynchus, Campylobacterota in Cx. pipiens, and Fusobacteriota in Ae. albopictus), consistent with global mosquito microbiome trends [15,16]. Genus- and species-level analyses further identified pathogenic or opportunistic taxa (e.g., Enterobacter, Campylobacter, Staphylococcus), underscoring their potential role in human infections under specific conditions.

Notably, species-specific abundance patterns were observed for key bacteria with public health implications. Wolbachia (clade B), exclusively detected in Ae. albopictus, has been shown to suppress dengue virus replication in Aedes aegypti by enhancing host immune responses [17] and reducing West Nile virus titers in cell lines [18]. Field-collected Ae. albopictus in Chengdu exhibited dual infections of Wolbachia wAlbB and an unclassified strain, diverging from the common wAlbA/wAlbB co-infections reported in Henan, and Nanjing [19,20]. This unique infection pattern warrants further investigation into its mechanistic and epidemiological implications. Similarly, Elizabethkingia anophelis, enriched in Cx. tritaeniorhynchus, has demonstrated inhibitory effects on Zika virus infection in Ae. aegypti [21] and Plasmodium development in Anopheles gambiae [22], suggesting its potential utility in biocontrol strategies.

Of particular concern is the high abundance of Klebsiella variicola and Klebsiella pneumoniae in Armigeres subalbatus. K. variicola, an emerging opportunistic pathogen, is associated with neonatal sepsis, pneumonia, and bloodstream infections (BSI), often exhibiting higher mortality rates than K. pneumoniae [23]. Although direct evidence of mosquito-to-human transmission remains lacking, the prevalence of these pathogens in mosquitoes highlights a potential public health risk.

Furthermore, phylogenetic analysis revealed unclassified taxa(*norank_d Bacteria,norank_p Candidatus_Hydrogenedentes*) clustering near Entomospira nematocera, a recently discovered mosquito-associated spirochete. These understudied lineages may represent novel functional microbiota, necessitating targeted cultivation or metagenomic approaches to elucidate their evolutionary origins and roles in mosquito biology.

KEGG pathway analysis indicated that Ae. albopictus microbiota were enriched in human disease-associated and genetic information processing pathways but depleted in cellular and environmental adaptation pathways compared to other species. These functional disparities may reflect evolutionary adaptations to ecological pressures or vector competence. While symbiotic bacteria are known to influence mosquito metabolism, immunity, and insecticide resistance, Chengdu-specific baseline data and mechanistic insights remain scarce. Future studies should integrate metagenomics, artificial symbiont colonization, and in vivo functional assays to validate these interactions and develop targeted vector control strategies.

